## Supplementary Material for "Distinct roles of dopamine and acetylcholine in delay- and effort-based decision making in humans"

***Supplementary Results*** ... 2

**1. Baseline Session Analysis** ... 2

**2. Model Validation and Parameter Recovery** ... 2

**3. Drug Effects on Vital Signs, Mood, Trail-Making Performance, MVC, and Effort Rating** ... 2

**4. Drug Effects on Decision Times** ... 4

**5. Computational Model Parameters & Self Ratings** ... 4

***Supplementary Methods*** ... 6

**6. Fitting Procedure Bayesian Regression** ... 6

**7. Fitting Procedure Hierarchical Bayesian Model** ... 7

7.1. Model Fitting and Comparison ... 7

7.2. Model Parametrization and Priors ... 9

**8. Physiological Measures and Bond-Lader Visual Analogue Scale** ... 10

**9. Decision Times Analysis** ... 11

**10. Computational Parameter Estimates, Self-Rating Scores, and Demographics** ... 11

***Supplementary References*** ... 13

***Supplementary Figures*** ... 15

***Supplementary Tables*** ... 24

### Supplementary Results

#### 1. Baseline Session Analysis

To analyse choice behaviour in the absence of drug manipulations, we conducted Logistic Bayesian Generalized Linear Mixed Models exclusively focusing on the data from the placebo (baseline) session. This analysis confirmed robust main effects of each task parameters on choice behaviour, demonstrating the expected patterns of reward and cost sensitivity across both tasks. Specifically, in both tasks, increased reward magnitudes were associated with a higher likelihood of choosing the high-cost option, while increased effort or delay levels decreased this likelihood (Supplementary Fig. 1 and Supplementary Table 4 and 5).

#### 2. Model Validation and Parameter Recovery

The simulated datasets mirror the original data reasonably well, confirming the model's ability to capture essential behavioural patterns observed in the actual data (Supplementary Fig. 2). Moreover, the parameter recovery revealed successful recovery of all group-level parameters, such that the mean of the simulated group-level parameters fell within the 95% HDI of the true parameter distribution (Supplementary Fig. 3). Similarly, averaged simulated and actual subject-level parameters were strongly correlated (all $r > 0.8$), indicating reliable estimation at the individual subject level (Supplementary Fig. 4).

#### 3. Drug Effects on Vital Signs, Mood, Trail-Making Performance, MVC, and Effort Rating

Bayesian Linear Mixed Models revealed that biperiden administration led to a reduction in heart rate and systolic blood pressure at both $T_1$ (heart rate: $HDI_{Mean}$ = -6.82, $HDI_{95\%}$ = [-10.74; - 2.97]; systolic blood pressure: $HDI_{Mean}$ = -3.91, $HDI_{95\%}$ = [-7.74; -0.13]) and $T_2$ (heart rate: $HDI_{Mean}$ = -7.88, $HDI_{95\%}$ = [-11.83; -3.89]; systolic blood pressure: $HDI_{Mean}$ = -4.80, $HDI_{95\%}$ = [-8.74; -0.95]), with no credible impact on diastolic blood pressure (Supplementary Fig. 5). Moreover, subjective ratings of alertness were credibly lower under both haloperidol and biperiden at $T_2$ (biperiden: $HDI_{Mean}$ = -0.15, $HDI_{95\%}$ = [-0.26; - 0.05]; haloperidol: $HDI_{Mean}$ = -0.15, $HDI_{95\%}$ = [-0.25; -0.04]; Supplementary Fig. 6). Notably, we did not find credible drug effects on the response times in the trail-making test A, the subjective rating of effort demand, and the MVC in the effort discounting task (Supplementary Fig. 7).

To investigate whether changes in physiological measures and mood ratings could be attributed to drug-induced changes in behaviour, we performed Bayesian correlation tests between each shift parameter that was credibly modulated by drug administration (i.e., $s\kappa_{HAL}$ and $s\beta_{BIP}$ in the delay discounting task and $s\kappa_{HAL}$, $s\kappa_{BIP}$, $s\beta_{HAL}$, and $s\beta_{BIP}$ in the effort discounting task) and the relative change in each control measure that credibly altered by drug intake (i.e., heart rate, systolic blood pressure, alertness ratings under biperiden and alertness ratings under haloperidol).

We found no credible correlation between any computational parameter that was modulated by biperiden and changes in systolic blood pressure at T1 in the effort ($r$ = 0.10, $HDI_{95\%}$ = [-0.14; 0.33] for $s\kappa_{BIP}$; $r$ = -0.16, $HDI_{95\%}$ = [-0.39; 0.07] for $s\beta_{BIP}$) and delay ($r$ = 0.06, $HDI_{95\%}$ = [-0.19; 0.29] for $s\beta_{BIP}$) discounting task. Similarly, we did not find any credible correlation between the computational parameters and biperiden-induced changes in systolic blood pressure at T2 ($r$ = -0.04, $HDI_{95\%}$ = [-0.27; 0.20] for $s\kappa_{BIP}$ (effort); $r$ = -0.08, $HDI_{95\%}$ = [-0.31; 0.17] for $s\beta_{BIP}$ (effort); $r$ = 0.13, $HDI_{95\%}$ = [-0.10; 0.35] for $s\kappa_{BIP}$ (delay)). The same applies for biperiden-induced reductions in heart rate at T1 ($r$ = 0.05, $HDI_{95\%}$ = [-0.18; 0.28] for $s\kappa_{BIP}$ (effort); $r$ = 0.06, $HDI_{95\%}$ = [-0.18; 0.28] for $s\beta_{BIP}$ (effort); $r$ = -0.06, $HDI_{95\%}$ = [-0.29; 0.18] for $s\kappa_{BIP}$ (delay)) and at T2 ($r$ = 0.23, $HDI_{95\%}$ = [-0.00; 0.45] for $s\kappa_{BIP}$ (effort); $r$ = -0.09, $HDI_{95\%}$ = [-0.32; 0.16] for $s\beta_{BIP}$ (effort); $r$ = -0.18, $HDI_{95\%}$ = [-0.40; 0.07] for $s\kappa_{BIP}$ (delay)) for both experimental paradigms. Alertness, which was affected by both drugs at T2, also did not show any credible correlations with biperiden- ($r$ = 0.14, $HDI_{95\%}$ = [-0.10; 0.37] for $s\kappa_{BIP}$ (effort); $r$ = 0.77, $HDI_{95\%}$ = [-0.16; 0.30] for $s\beta_{BIP}$ (effort), $r$ = -0.16, $HDI_{95\%}$ = [-0.38; 0.08] for $s\beta_{BIP}$ (delay)) as well as haloperidol-induced changes ($r$ = -0.05, $HDI_{95\%}$ = [-0.28; 0.19] for $s\kappa_{HAL}$ (effort); $r$ = -0.09, $HDI_{95\%}$ = [-0.33; 0.16] for $s\beta_{HAL}$ (effort); $r$ = 0.05, $HDI_{95\%}$ = [-0.20; 0.29] for $s\kappa_{HAL}$ (delay)).

### 4. Drug Effects on Decision Times

Having established distinct task-specific drug effects on participants' choice behaviour, we next asked how the drugs affected the dynamics of choice, as reflected in how decision times were modulated by key decision variables. We investigated this with Bayesian Linear Mixed Models, using log-transformed decision times on each trial as the dependent variable. In both tasks, higher reward magnitudes decreased, while higher cost levels (effort or delay) increased decision times (reward effect on effort discounting: $HDI_{Mean}$ = -0.183, $HDI_{95\%}$ = [-0.213; -0.154]; reward effect on delay discounting: $HDI_{Mean}$ = -0.168, $HDI_{95\%}$ = [-0.190; -0.145]; effort effect on effort discounting: $HDI_{Mean}$ = 0.084, $HDI_{95\%}$ = [0.063; 0.105]; delay effect on delay discounting: $HDI_{Mean}$ = 0.036, $HDI_{95\%}$ = [0.020; 0.053]; Supplementary Fig. 8b-c). Notably, in both tasks, haloperidol attenuated this speeding effect of reward magnitude, indexed by a credible interaction effect between haloperidol and reward in both tasks (effort discounting: $HDI_{Mean}$ = 0.036, $HDI_{95\%}$ = [0.009; 0.063]; delay discounting: $HDI_{Mean}$ = 0.029, $HDI_{95\%}$ = [0.011; 0.046]: Supplementary Fig. 9b, 9e). Haloperidol further attenuated the decelerating effect of delay ($HDI_{Mean}$ = -0.020, $HDI_{95\%}$ = [-0.036; -0.003]; Fig. 9f), but not effort ($HDI_{Mean}$ = -0.021, $HDI_{95\%}$ = [-0.047; 0.003]; Supplementary Fig. 9c). This aligns with the decreased delay sensitivity observed in the delay discounting task analysis. Moreover, haloperidol administration induced an overall increase in decision times in the delay discounting task ($HDI_{Mean}$ = -0.077, $HDI_{95\%}$ = [-0.123; -0.031]; Supplementary Fig. 9d), while this effect was not observed in the effort discounting task ($HDI_{Mean}$ = 0.016, $HDI_{95\%}$ = [-0.047; 0.078]; Supplementary Fig. 9a). Unlike under haloperidol, none of the effects of task parameters on response speed were modulated by biperiden. See Supplementary Table 6 and 7 for full results.

### 5. Computational Model Parameters & Self Ratings

Robust linear regression models did not reveal any significant associations between sex, self-reported questionnaire ratings (total scores and subscales), and both effort and delay baseline (placebo-condition) discounting parameters (Supplementary Tables 8 – 11). However, the analysis did reveal a significant main effect of individuals age on the effort discounting parameter, indicating a higher tendency to discount rewards in older compared to younger participants (*beta* = 0.007, $p$ = 0.008).

### Supplementary Methods

6. Fitting Procedure Bayesian Regression

To examine the impact of varying levels of reward and/or costs, as well as the administered drug, on choosing the high-reward/high-cost option, we employed Logistic Bayesian Generalized Linear Mixed Models. For the effort discounting task, the fixed effects included reward (difference between reward magnitudes of the high-cost versus low-cost option), effort (difference between effort requirement of the high-cost versus low-cost option), the administered drug (with placebo set as the reference category), and their interactions. For the delay discounting task, we followed a similar approach, but instead of using difference values, we used the absolute reward and delay levels of the high-cost option, given that the low-cost option remains fixed to a constant value throughout the task. Furthermore, to mitigate the risk of false positive results, all models contained a full random-effects structure (Barr et al., 2013). This approach allowed us to account for individual variability, leading to more robust and reliable findings.

To control for potential confounding effects of fatigue and session, we ran additional separate GLMMs, extending the fixed-effects structures. Specifically, to test for fatigue, we included trial number and its two-way interactions with drug as additional predictors. In separate models, we controlled for session effects by adding session, as well as the two-way interactions between session and drug.

Further, to gain insights into choice behaviour in the absence of any pharmacological manipulations, we conducted additional GLMMs exclusively analysing data from the placebo sessions. These models were identical to the previously described regression analyses, including the full random-effects structure, but excluded the *drug* predictor. This analysis provided estimates of the main effects of task manipulations (i.e., reward and cost sensitivity) under the baseline condition.

To ensure robust and informative Bayesian parameter estimation and avoid issues of unstable parameter estimation that could appear with noninformative and flat priors, we followed the approach recommended by Gelman et al. and implemented weakly informative priors (2008). First, we standardized all nonbinary variables to have a mean of 0 and a standard deviation of 0.5. Then, we used the following priors in our analyses:

| | |
|---|---|
| Regression Coefficient | $\beta \sim Cauchy(0, 2.5)$ |
| Intercept | $\beta_0 \sim Cauchy(0, 10)$ |
| Correlation Matrix | $\Sigma \sim LKJcorr(1)$ |
| Standard Deviation | $\sigma \sim Student(3, 0, 2.5)$ |

### 7. Fitting Procedure Hierarchical Bayesian Model

#### 7.1. Model Fitting and Comparison

To identify the model that best describes how rewards are devalued by increasing levels of effort and delay, we employed four commonly used discounting models on participants' choice data from both tasks (Białaszek et al., 2017; Chong et al., 2017; Hartmann et al., 2013; Klein-Flügge et al., 2015): linear (Supplementary Eq. 1), parabolic (Supplementary Eq. 2), hyperbolic (Supplementary Eq. 3), and exponential (Supplementary Eq. 4). To minimize the potential impact of drug administration on the discounting function, we restricted the model fitting to choice data from the (baseline) placebo condition.

$$SV(t) = R(t) - \kappa * C(t) \tag{1}$$

$$SV(t) = R(t) - \kappa * C(t)^2 \tag{2}$$

$$SV(t) = \frac{R(t)}{1 + \exp(\kappa) * C(t)} \tag{3}$$

$$SV(t) = R(t) * e^{-\kappa * C(t)} \tag{4}$$

All discounting models assume that the subjective value (SV) of an offer is calculated by taking into account the reward (R) and the cost (C) level on trial (t). In the effort discounting task, the cost level is represented by the effort level, which is scaled to the proportion of the maximum voluntary contraction. In the delay discounting task, the cost level is represented by the delay of the high-cost option, indicating the number of days necessary to wait to obtain the reward. The degree to which rewards are discounted by increasing levels of costs is modelled by a subject-specific discounting parameter ($\kappa$), which quantifies the steepness of each individual's devaluation of rewards as costs increase. Higher values of $\kappa$ represent higher steepness in devaluation, indicating stronger sensitivity to increasing costs, while lower values represent lower steepness, indicating less sensitivity to increasing costs. Importantly, in the effort discounting task, two options with varying levels of reward and effort are presented on each trial, leading to the calculation of two different SVs. In contrast, in the delay discounting task, one option varies in reward and delay, while the other SV is fixed at 20, resulting in the calculation of a single SV for that option per trial. Additionally, note that the $\kappa$ values for the delay discounting task were modelled in log space to prevent numerical instability caused by highly skewed $\kappa$ values. SVs for the high-reward/high-cost (HC) and the low-reward/low-cost (LC) were then transformed to choice probabilities, using a softmax function (Eq. 5).

$$P\left(HC_{(t)}\right) = \frac{\exp\left(SV(HC_{(t)}) * \beta\right)}{\exp\left(SV(HC_{(t)}) * \beta\right) + \exp\left(SV(LC_{(t)}) * \beta\right)} \quad (5)$$

The choice consistency was modelled using the inverse temperature parameter $\beta$.

To determine the best fitting models for describing participants' behaviour in each task, we employed the leave-one-out information criterion (LOOIC), which estimates out-of-sample prediction accuracy by utilizing the log-likelihood. The LOOIC was assessed using the loo package in R (Vehtari et al., 2017). Lower LOOIC values indicate better model fit, similar to traditional information criteria such as AIC and BIC (see Supplementary Table 3).

### 7.2. Model Parametrization and Priors

For all hierarchical models, we assume that the subject-level parameters are drawn from group-level normal distributions. We use Uniform and half-Cauchy distributions for the group-level mean ($\mu$) and standard deviations ($\sigma$) of the (baseline) placebo-condition discounting parameters $\kappa$ and inverse temperature $\beta$, respectively. For all shift parameters, which indicate drug-specific effects on $\kappa$ and $\beta$, Gaussian prior distributions were used for the means, and half-Cauchy distributions were used for the standard deviations. Based on previous findings, the standard deviations of all half-Cauchy distributions were set with a location of 0 and a scale of 2.5. Additionally, more restrictive priors were set for the Gaussian distribution of all group-level shift hyperparameters with a location of 0 and a scale of 2 (Knauth & Peters, 2022; Mathar et al., 2022; Wagner et al., 2020). To account for the different degrees of discounting depending on the cost type and the logarithmic transformation of $\kappa$ in delay discounting, we used distinct ranges for the Uniform distribution of the group-level means in both tasks. These ranges were based on numerically plausible values and previous findings (Knauth & Peters, 2022; Lockwood et al., 2021, 2022; Mathar et al., 2022; Wagner et al., 2020).

In summary, the prior distributions for our hierarchical models are as follows:

$$\mu_{\kappa(Effort)} \sim Uniform(0, 5)$$

$$\mu_{\kappa(Delay)} \sim Uniform(-20, 3)$$

$$\sigma_{\kappa} \sim HalfCauchy(0, 2.5)$$

$$\mu_{\beta} \sim Uniform(0, 10)$$

$$\sigma_{\beta} \sim HalfCauchy(0, 2.5)$$

$$\mu_{S_x} \sim Normal(0, 2)$$

$$\sigma_{S_x} \sim HalfCauchy(0, 2.5)$$

8. Physiological Measures and Bond-Lader Visual Analogue Scale

Participants completed subjective mood ratings using the German version of the Bond and Lader visual analogue scale (Bond & Lader, 1974) at three different time points ($T_0$, $T_1$, and $T_2$). Additionally, blood pressure and heart rate measurements were taken at the same time points using a digital blood pressure monitor (OMRON model M500, Healthcare Europe B.V., The Netherlands). These manipulation checks were conducted at $T_0$ (before drug administration), $T_1$ (before starting the task, approximately 170 min after haloperidol or 50 min after biperiden intake), and $T_2$ (after finishing the task, approximately 230 min after haloperidol or 110 min after biperiden intake). These assessments allowed us to examine potential effects of the administered drugs on subjective mood states and physiological parameters. Subjective mood rating scales involved 16 binary items presented on a horizontal line on a sheet of paper. Each item consisted of two words describing opposing mood states (e.g., "happy versus sad"), and participants indicated their mood by marking the line closer to one of the two words. Based on a factor analysis using a principal component solution and orthogonal rotation of the factor matrix, three separate factor scores were extracted: alertness, contentedness, and calmness. Self-ratings were analysed by measuring the distance in millimetres from the end of the line to the subject's mark. These measurements were then log-transformed to correct for skewness (Bond & Lader, 1974). In addition, once per session, participants completed an effort rating, in which they were required to rate each effort level they encountered during the tasks. Furthermore, prior to beginning the effort discounting task, participants completed the trail-making test A, and we measured participants' maximum voluntary contraction (MVC).

To analyse the effects of the drugs on mood ratings, physiological measures, trail-making test response times, MVC, and subjective effort perception ratings, we applied Bayesian Linear Mixed Effects Models. For mood ratings and physiological measures, the models included the factors time ($T_0$, $T_1$, and $T_2$), drug (PLC, HAL, BIP), and their interaction as fixed effects, with subject-specific intercepts. Similarly, for the trail-making test, MVC, and effort ratings, which were measured once per session, the models included drug (PLC, HAL, BIP), session (Session 1, Session 2, Session 3), and their interaction as fixed effects, with subject-specific intercepts. For analysing effort ratings, we additionally included effort levels as fixed effects in the model. For parameter estimation, we used non-informative priors (the brms default) and ran four chains with 3000 samples (1000 samples for warmup).

Importantly, to test whether credible changes in physiological parameters or mood ratings induced by drug administration could explain changes in behaviour, we used Bayesian correlation tests to examine possible associations. We correlated difference values ($T_0$ vs. timepoints with credible drug-induced influences; i.e., $T_2$ for mood ratings, $T_1$ and $T_2$ for physiological parameters) with the mean estimates of all shift parameters that were credibly modulated by either haloperidol or biperiden.

### 9. Decision Times Analysis

Similarly, to the model-agnostic analysis of the choice data, we performed a separate analysis focusing on participants' decision times in both tasks using Bayesian Linear Mixed Models. In contrast to the previous analysis, with binary choice data as the outcome variable with a Bernoulli response distribution and a logit link function, here, we used the log-transformed decision times (in milliseconds) as the outcome variable with a Gaussian distribution function. We then regressed the decision times to the same set of fixed-effect predictors, including drug, reward, cost (i.e., delay or effort), and their respective interaction terms.

However, in order to ensure full model convergence, we reduced the random-effects structure (the full random-effects structure led to convergence issues, indicated by r-hat values > 1.05). Specifically, we removed the three-way interaction effect (interaction between drug, reward, and cost type), including only main and two-way interaction effects in the random effects structure. As mentioned earlier, we applied weakly informative priors, scaling nonbinary variables to have a mean of 0 and a standard deviation of 0.5. Posterior distributions of parameter estimates were obtained by running four chains with 3000 samples, including 1000 samples for warmup.

### 10. Computational Parameter Estimates, Self-Rating Scores, and Demographics

As a last step of our analysis, we examined potential associations between the discounting parameters $\kappa$ and self-reported questionnaire ratings, including the Apathy Evaluation Scale (AES) and Barratt Impulsiveness Scale-15 (BIS-15), with all subscales, and Beck Depression Inventory (BDI). We also investigated the relationship of both model parameters with demographic variables (sex and age). To this aim, we conducted four separate robust linear regressions, using the mean estimates of the discounting parameter $\kappa$ from the placebo condition of both tasks, serving as a baseline value for each participants' discounting behaviour. We chose robust regression models because they have been shown to be less sensitive to the influence of outliers (Yu & Yao, 2017). We regressed this outcome against predictors of age, sex, and the z-scored total scores of all questionnaire scores. Further, to gain insights into the effects of each subscale of the apathy and impulsivity questionnaires, we again performed separate robust regression models, this time using the z-scored subscale scores of both questionnaires as predictors.

### Supplementary Figures

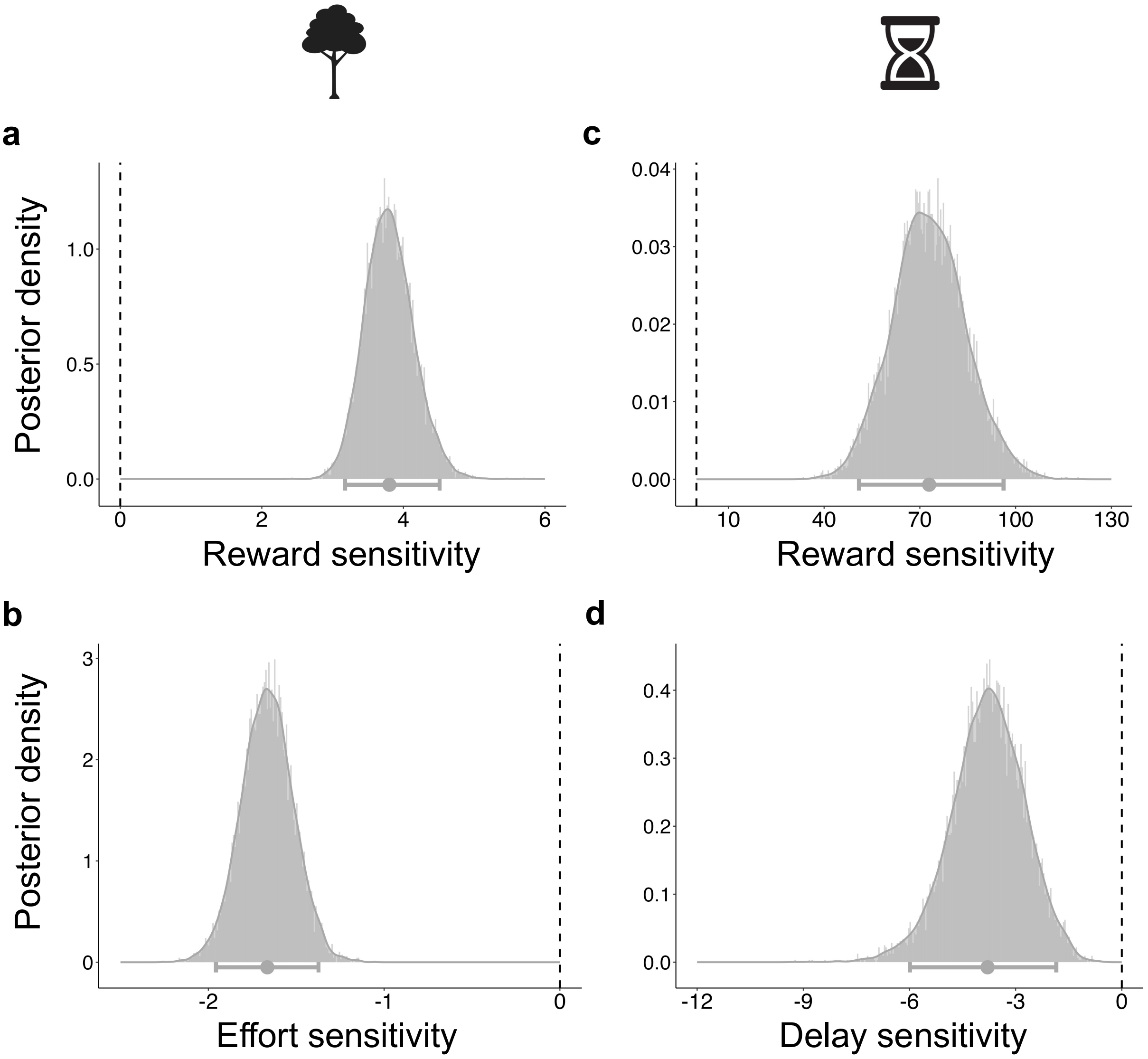


**Supplementary Fig. 1. Choice Behaviour in the placebo (baseline) condition.** Posterior distributions and 95% HDI of the Logistic Bayesian Generalized Linear Mixed Models depict the estimate of each task parameter on choosing the high-cost option. (**a**) Higher reward magnitudes increased the overall willingness to invest physical effort for a corresponding reward in the effort discounting task. (**b**) Higher levels of effort had the opposite effect. (**c**) Similarly, higher reward magnitudes increased the likelihood to choose the high-cost option in the delay discounting task. (**d**) In contrast, higher levels of delay decreased the willingness to choose the high-reward/high-delay option. Bold dots represent the mean group-level estimate of the posterior distribution. The horizontal bars represent the group-level 95% highest density interval.

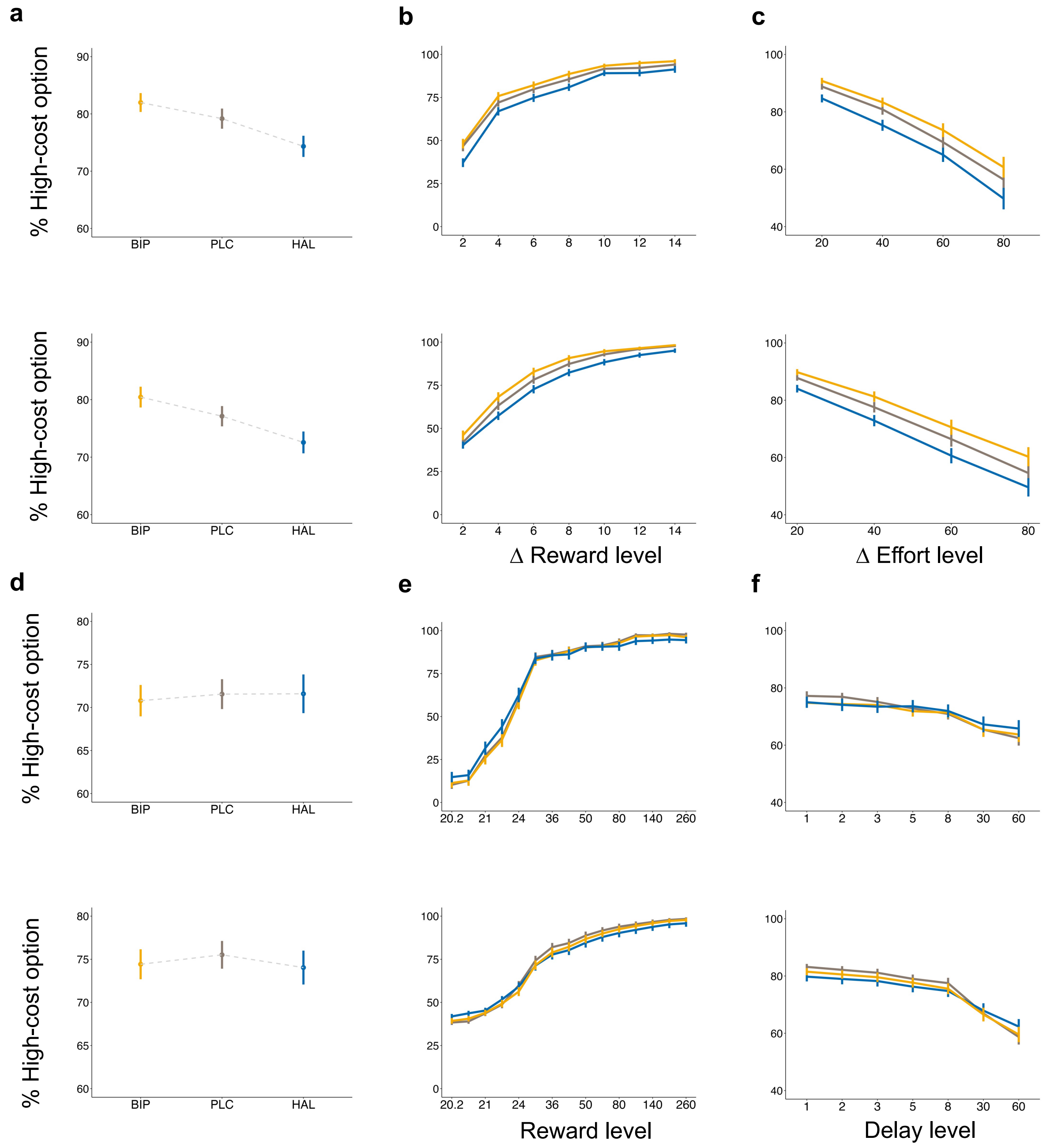


**Supplementary Fig. 2. Model Validation for the Effort (a-c) and Delay Discounting Task (d-f).** Plots depict the averaged overall proportion of choosing the high-cost option as a function of reward and cost for both the effort (**a, b, c**) and the delay (**d, e, f**) discounting task. The upper panels display the actual data and the lower panels present the simulated data for comparison. Group-level means are indicated by dots, with error bars representing the standard error of the mean. In the effort discounting task, reward levels are presented as the difference in magnitude between the high- and low-cost option, and in the delay discounting task, reward levels are shown as the reward value of the high-cost option. Likewise, the effort level corresponds to the difference between the proportions of the individually calibrated MVC of the high and low-cost option, while the delay levels indicate the delay of the high-cost option.

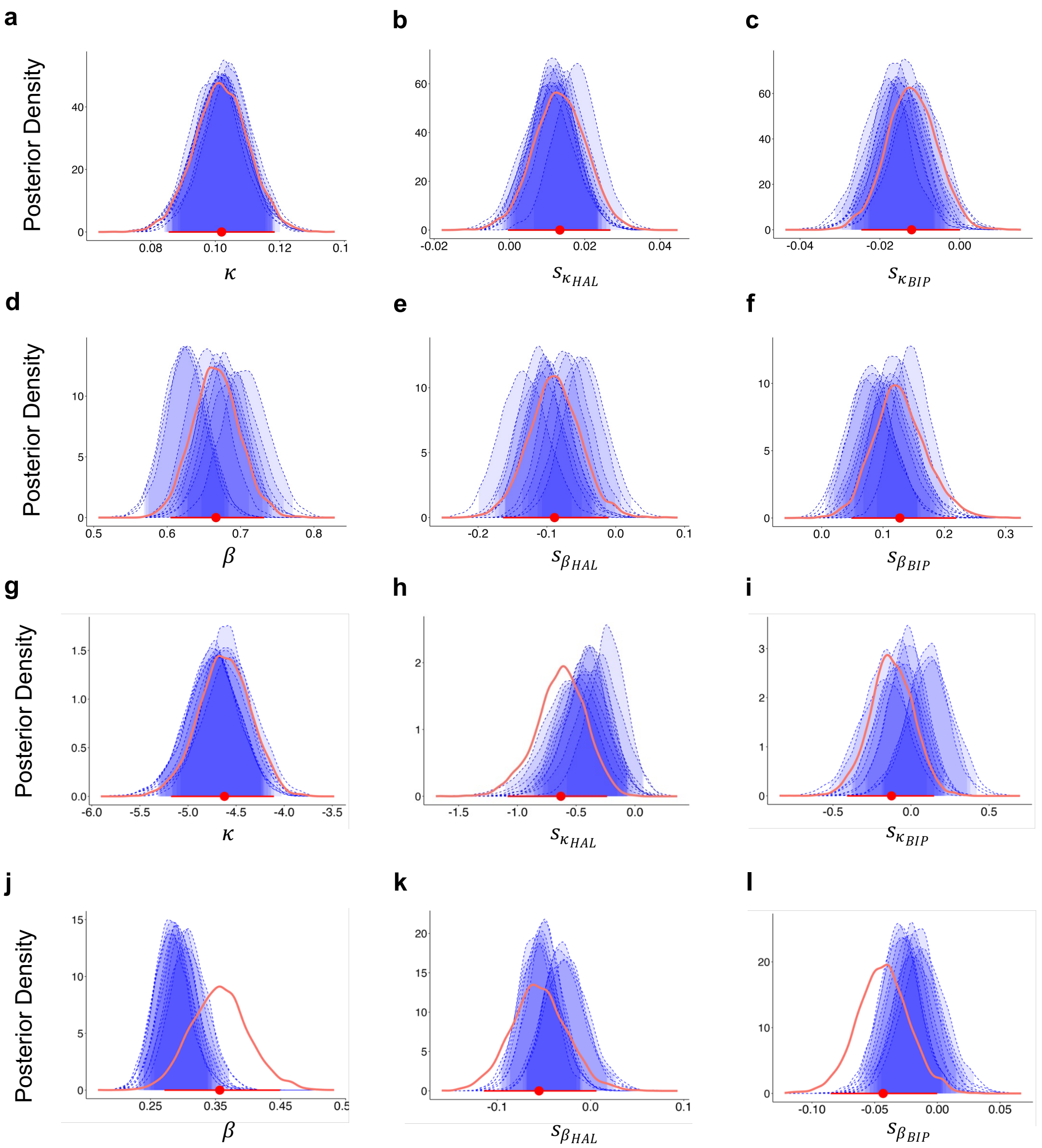


**Supplementary Fig. 3. Parameter Recovery of the Group-Level Posterior Distributions for the Effort (a-f) and Delay Discounting Task (g-l).** We tested the ability of our models to recover parameters using simulated datasets. Each panel displays the actual distribution (red) of each parameter of interest alongside ten corresponding simulated datasets (blue). Horizontal bars, depicted in red, represent the group-level 95% HDI of the actual parameter estimate, with dots indicating the mean of the distribution. The shaded area in blue depicts the 95% HDI of the simulated parameter estimates. We evaluated whether the mean estimates of the simulated group-level parameter values fell within the 95% HDI of the true parameter distribution. The parameter recovery analysis of the group-level distribution demonstrated positive results, as all mean parameter estimates of the simulated data are located within the 95% HDI of the actual dataset.

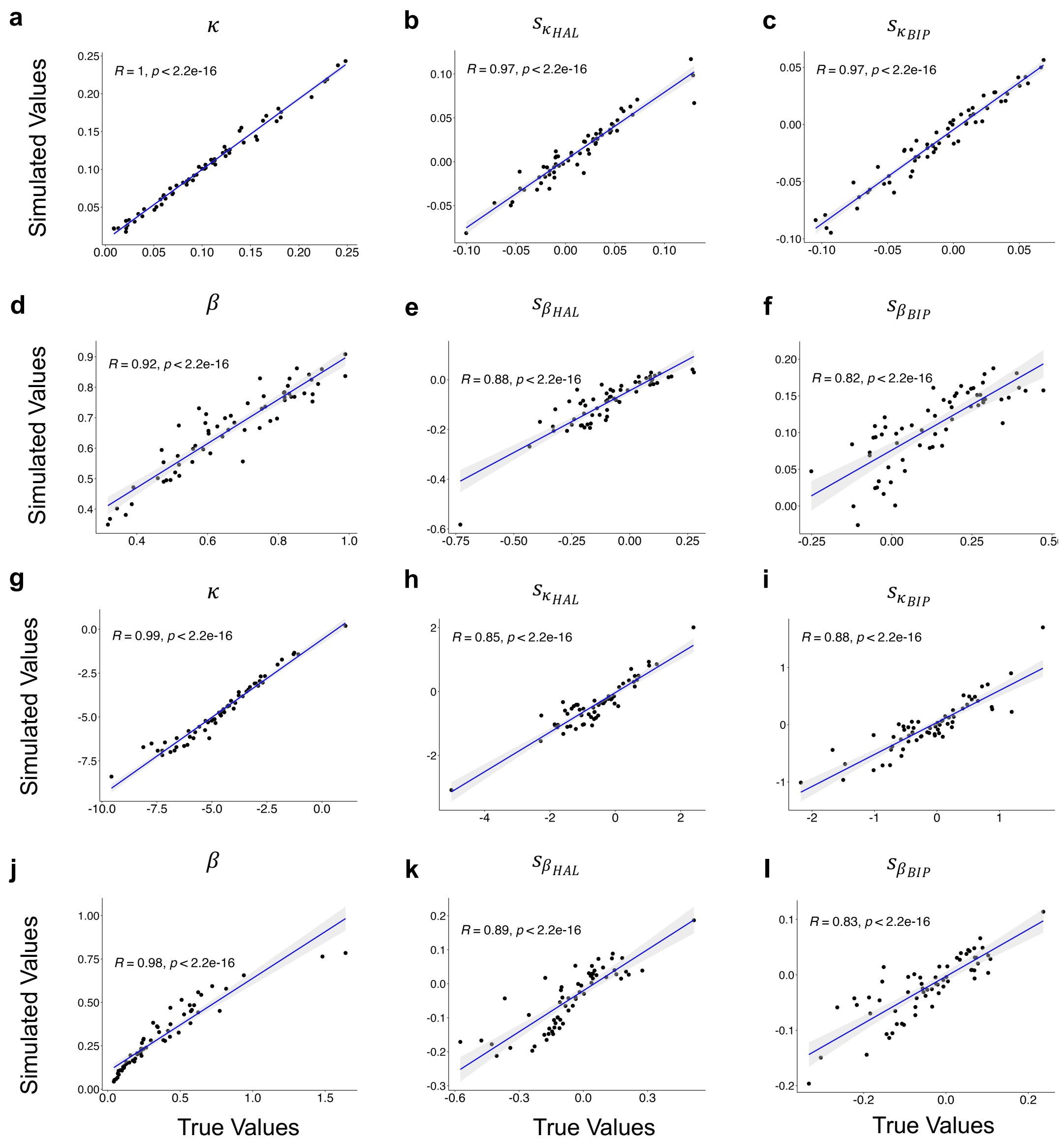


**Supplementary Fig. 4. Parameter Recovery of the Subject-Level Parameters for the Effort (a-f) and Delay Discounting Task (g-l).** We calculated the Pearson correlation coefficients between the mean subject-level posterior distribution of the simulated and actual data. For the simulated data, subject-level means were averaged across each dataset. The correlation coefficients demonstrate strong to excellent correlations (all $r > 0.8$), further confirming that the models are able to accurately to recover the actual task parameter values.

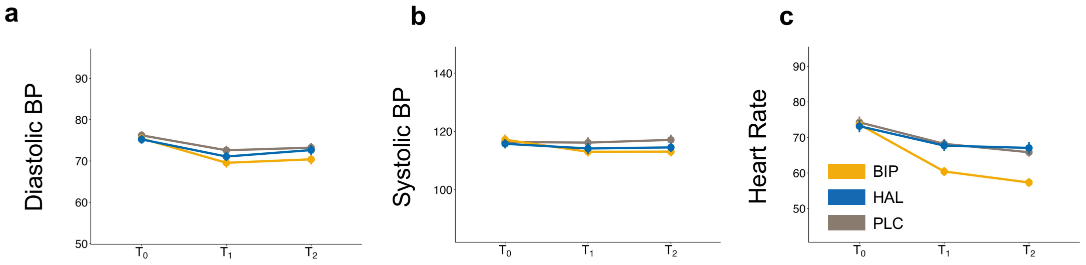


**Supplementary Fig. 5. Physiological Effects of Drug Administration.** Plots depict the change in each physiological parameter following drug administration. (**a**) Diastolic BP decreased at T1 ($HDI_{Mean}$ = -3.75, $HDI_{95\%}$ = [-5.71; -1.85]) and T2 ($HDI_{Mean}$ = -2.99, $HDI_{95\%}$ = [-4.96; -1.07]) compared to T0. However, we did not find a credible main or interaction effect of drug, implying that diastolic BP decreases as the task progresses irrespective of drug administration. (**b**) Systolic BP reflected notable two-way biperiden interactions at T1 and T2 (Biperiden x T1: $HDI_{Mean}$ = -3.91, $HDI_{95\%}$ = [-7.74; -1.07]; Biperiden x T2: $HDI_{Mean}$ = -4.80, $HDI_{95\%}$ = [-8.74; -0.95]), suggesting a more pronounced decrease in systolic BP following Biperiden application. (**c**) Heart rate was reduced at T1 ($HDI_{Mean}$ = -6.82, $HDI_{95\%}$ = [-10.74; -2.97]) and T2 ($HDI_{Mean}$ = -7.88, $HDI_{95\%}$ = [-11.83; -3.89]) relative to T0. A credible two-way interaction was found between Biperiden at T1 and T2 (Biperiden x T1: $HDI_{Mean}$ = -6.82, $HDI_{95\%}$ = [-10.74; -2.97]; Biperiden x T2: $HDI_{Mean}$ = -7.88, $HDI_{95\%}$ = [-11.83; -3.89]), indicating that, analogously to the drop in systolic BP, the drop in heart rate at T1 and T2 is more pronounced following biperiden administration compared to placebo. Dots represent the group-level mean, error bars depict the standard error of the mean.

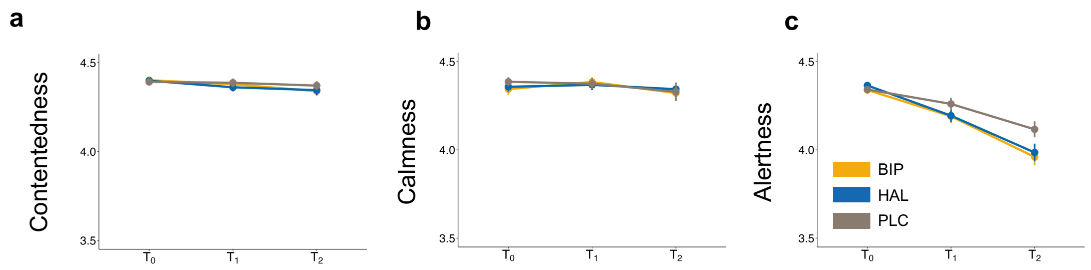


**Supplementary Fig. 6. Subjective Mood Rating Effects of Drug Administration.** Plots show alterations in subjective mood ratings following drug administration. No credible effects of time or drug were found for (**a**) contentedness and (**b**) calmness ratings. However, (**c**) alertness ratings notably decreased over time, exhibiting a credible drop at both T1 ($HDI_{Mean}$ = -0.08, $HDI_{95\%}$ = [-0.16; -0.01]) and T2 ($HDI_{Mean}$ = -0.23, $HDI_{95\%}$ = [-0.30; -0.15]) compared to T0. Importantly, at T2, credible two-way interaction effects were found for biperiden and haloperidol (Biperiden x T2: $HDI_{Mean}$ = -0.15, $HDI_{95\%}$ = [-0.26; -0.05]; Haloperidol x T2: $HDI_{Mean}$ = -0.15, $HDI_{95\%}$ = [-0.25; -0.04]), suggesting that both drugs led to more pronounced reductions in alertness ratings towards the end of the experiment, compared to placebo. Dots represent the group-level mean, error bars depict the standard error of the mean.

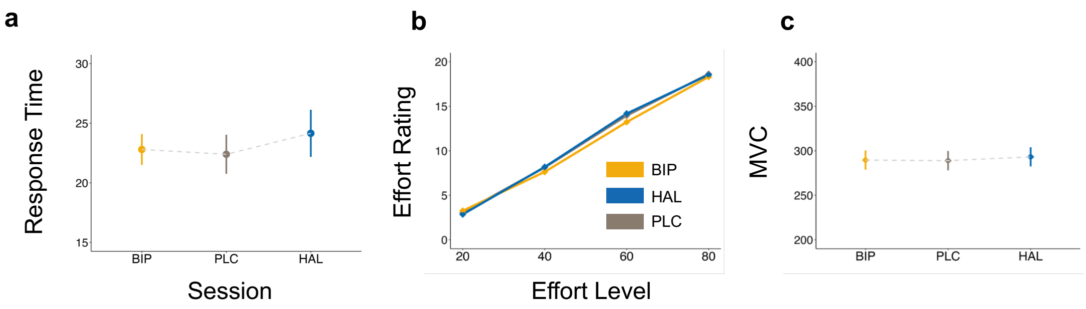


**Supplementary Fig. 7. Drug Effects on Trail-Making Test Performance, Subjective Effort Ratings, and MVC.** (**a**) No credible drug effects were observed for response times during trail-making test A. However, credible main effects of session were found (Session 2: $HDI_{Mean}$ = -4.77, $HDI_{95\%}$ = [-8.41; -1.17]; Session 3: $HDI_{Mean}$ = -6.40, $HDI_{95\%}$ = [-10.14; -2.66]), suggesting that participants became faster after the first session, possibly due to familiarity with the task. (**b**) Effort ratings were modulated only by increasing effort levels ($HDI_{Mean}$ = 5.15, $HDI_{95\%}$ = [4.81; 5.49]), suggesting no drug effects on the subjective experience of effort demand. (**c**) MVC was not credibly modulated by drug administration, indicating that participant's ability to exert effort was not modulated by drug either. Dots represent the group-level mean and error bars represent standard error of the mean.

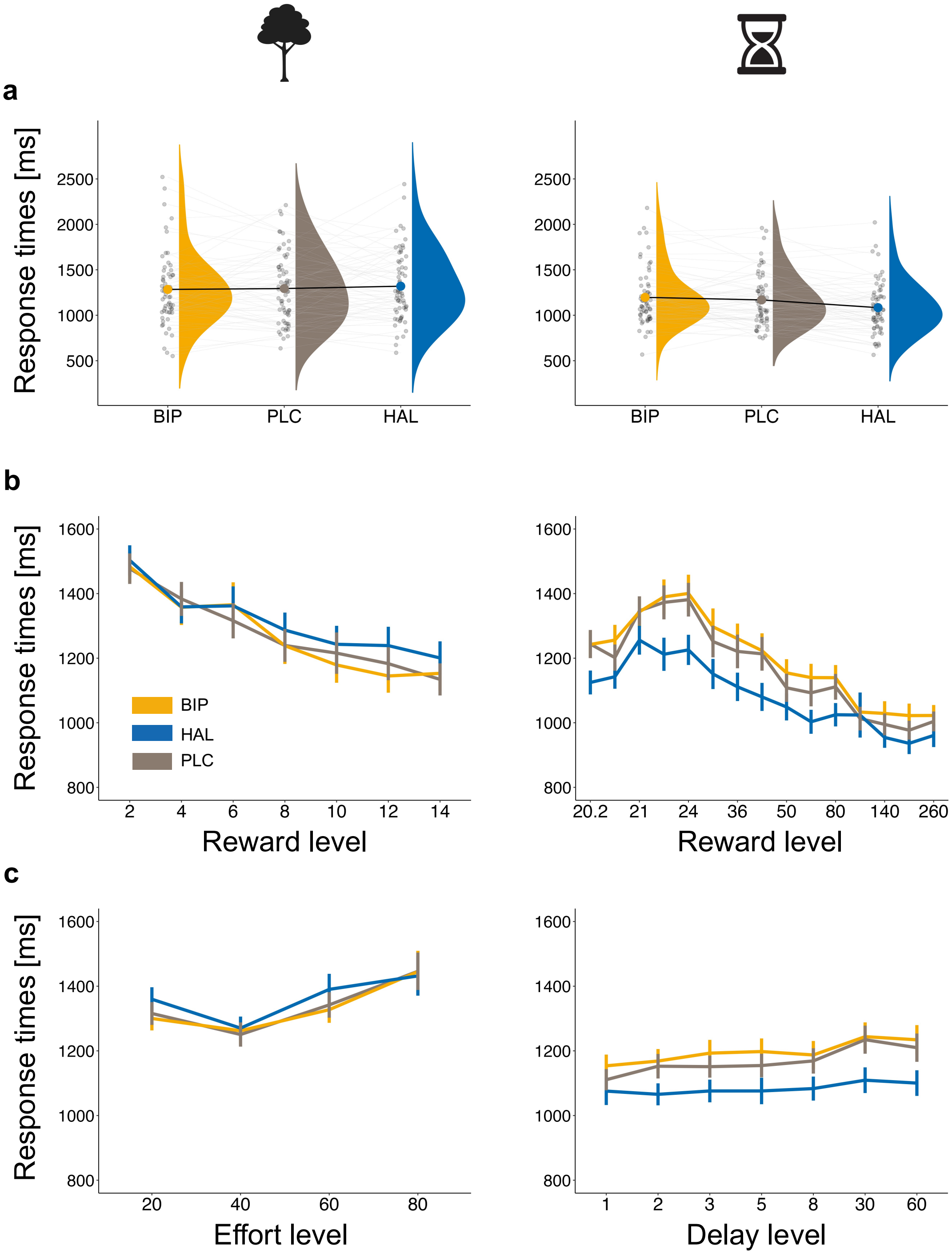


**Supplementary Fig. 8. Decision Times in milliseconds (ms) in the Effort (left panel) and Delay (right panel) Discounting Tasks**. (**a**) Overall decision times in the effort discounting task (left) were not affected by haloperidol or biperiden. However, in the delay discounting task (right), haloperidol reduced the decision times. (**b**) Decision times decreased with larger rewards. Haloperidol reduced this speed-up effect in both the effort (left) and delay discounting task (right panel). (**c**) Conversely, decision times increased with higher cost levels. This effect was not modulated by any drug in the effort discounting task (left). However, in the delay discounting task (right), haloperidol diminished the decelerating effect of increasing delay levels. **a** shows group-level (single-subject) means represented by bold (light) dots. **b** and **c** display averaged group-level means per reward and cost level, with error bars representing the standard error of the mean. Reward levels are presented as the difference in magnitude between the high- and low-cost option in the effort discontinuing task and as the absolute reward value of the high-cost option in the delay discounting task. Likewise, the effort level represents the difference between the proportions of the individually calibrated MVC of the high- and low-cost option, while the delay level indicates the delay of the high-cost option.

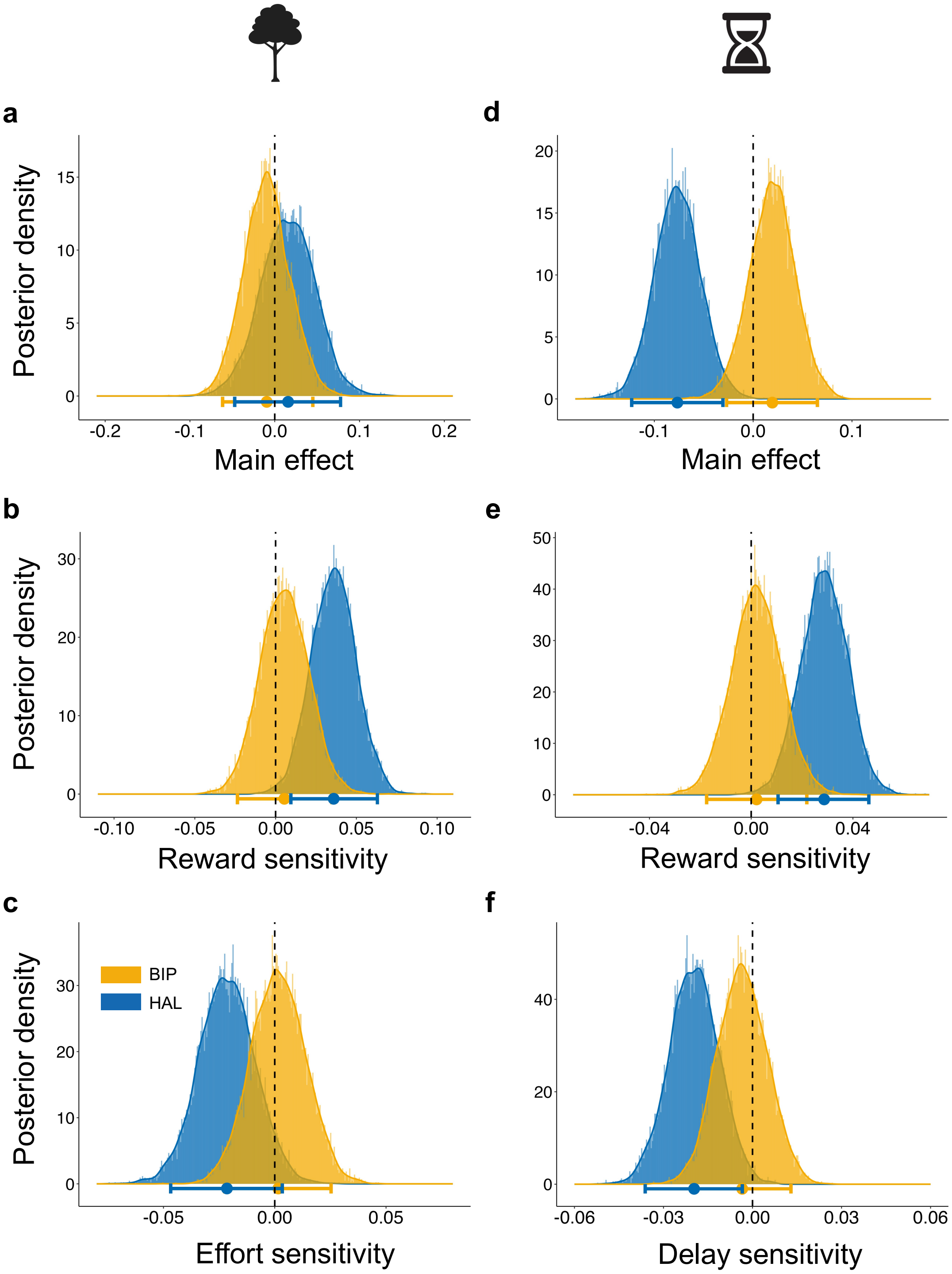


**Supplementary Fig. 9. Drug Effects on Decision Times.** Posterior distributions and 95% HDI of the Bayesian Linear Mixed Models depict the estimate of each effect on decision times. (**a**) In the effort discounting task, overall decision times were not credibly influenced by either haloperidol or biperiden. (**b**) Notably, a haloperidol-by-reward interaction revealed a reduced reward sensitivity after haloperidol administration. (**c**) On the other hand, sensitivity towards increasing levels of effort is not affected by either drug. (**d**) In the delay discounting task, haloperidol credibly reduced overall decision times, while biperiden did not show any credible effect. (**e**) Analogous to the effort discounting task, a credible haloperidol-by-reward interaction demonstrate reduced reward sensitivity following haloperidol administration. (**f**) Furthermore, haloperidol administration credibly reduced delay sensitivity, while biperiden had no effect on the impact of delay on decision times. Bold dots represent the mean group-level estimate of the posterior distribution. The horizontal bars represent the group-level 95% highest density interval.

### Supplementary Tables

**Supplementary Table 1.** Bayesian Generalized Linear Mixed Models of the Effort Discounting Task, Regressing Choices (High-Cost vs. Low-Cost Option) on Predictors for Drug, Reward (Difference between High-Cost vs. Low-Cost Reward Level), Effort (Difference between High-Cost vs. Low-Cost Effort Level), and their Interaction Terms.

| **Parameter** | **Estimate** | **Est. Error** | **2.5%** | **97.5%** |
|---|---|---|---|---|
| **(Intercept)** | 2.544 | 0.219 | 2.116 | 2.988 |
| **Biperiden** | 0.620 | 0.207 | 0.230 | 1.048 |
| **Haloperidol** | -0.532 | 0.203 | -0.943 | -0.136 |
| **Reward** | 3.436 | 0.251 | 2.961 | 3.945 |
| **Effort** | -1.638 | 0.122 | -1.874 | -1.402 |
| **Biperiden x Reward** | 0.802 | 0.292 | 0.256 | 1.409 |
| **Haloperidol x Reward** | -0.296 | 0.257 | -0.797 | 0.203 |
| **Biperiden x Effort** | -0.011 | 0.153 | -0.309 | 0.303 |
| **Haloperidol x Effort** | 0.084 | 0.121 | -0.150 | 0.321 |
| **Reward x Effort** | 0.150 | 0.183 | -0.212 | 0.509 |
| **Biperiden x Reward x Effort** | 0.104 | 0.295 | -0.476 | 0.675 |
| **Haloperidol x Reward x Effort** | -0.223 | 0.238 | -0.682 | 0.256 |

**Supplementary Table 2.** Bayesian Generalized Linear Mixed Models of the Delay Discounting Task, Regressing Choices (High-Cost vs. Low-Cost Option) on Predictors for Drug, Reward (High-Cost Option Reward), Delay (High-Cost Option Delay), and their Interaction Terms.

| **Parameter** | **Estimate** | **Est. Error** | **2.5%** | **97.5%** |
|---|---|---|---|---|
| **(Intercept)** | 19.555 | 2.629 | 14.436 | 24.643 |
| **Biperiden** | -1.315 | 1.263 | -4.147 | 0.655 |
| **Haloperidol** | -0.046 | 1.095 | -2.648 | 1.884 |
| **Reward** | 55.637 | 7.095 | 41.762 | 69.382 |
| **Delay** | -2.291 | 0.522 | -3.303 | -1.250 |
| **Biperiden x Reward** | -3.871 | 3.478 | -11.873 | 1.453 |
| **Haloperidol x Reward** | -1.271 | 2.952 | -8.257 | 4.037 |
| **Biperiden x Delay** | 0.781 | 0.481 | -0.104 | 1.807 |
| **Haloperidol x Delay** | 1.332 | 0.540 | 0.328 | 2.440 |
| **Reward x Delay** | -2.804 | 1.382 | -5.521 | -0.142 |
| **Biperiden x Reward x Delay** | 1.054 | 1.333 | -1.361 | 4.003 |
| **Haloperidol x Reward x Delay** | 2.383 | 1.525 | -0.465 | 5.514 |

**Supplementary Table 3.** Model comparison for the effort and delay discounting task. To compare the validity of each model, we used the leave-one-out cross-validation information criterion (LOOIC) procedure. A lower LOOIC score indicates a better-fitting model.

| **Model** | ***Effort*** | ***Delay*** |
|---|---|---|
| | **LOOIC** | **LOOIC** |
| *Parabolic* | 15109.7* | 27279.6 |
| *Linear* | 17337.5 | 26041.5 |
| *Hyperbolic* | 18477.9 | 25662.4* |
| *Exponential* | 17961.2 | 26217.5 |

**Supplementary Table 4.** Bayesian Generalized Linear Mixed Models of the Effort Discounting Task – Baseline Session; Regressing Choices (High-Cost vs. Low-Cost Option) on Predictors for Reward (Difference between High-Cost vs. Low-Cost Reward Level), Effort (Difference between High-Cost vs. Low-Cost Effort Level), and their Interaction Terms.

| Parameter | Estimate | Est. Error | 2.5% | 97.5% |
|---|---|---|---|---|
| (Intercept) | 2.747 | 0.288 | 2.199 | 3.331 |
| Reward | 3.802 | 0.342 | 3.174 | 4.512 |
| Effort | -1.664 | 0.148 | -1.957 | -1.373 |
| Reward x Effort | 0.120 | 0.235 | -0.345 | 0.587 |

**Supplementary Table 5.** Bayesian Generalized Linear Mixed Models of the Delay Discounting Task – Baseline Session; Regressing Choices (High-Cost vs. Low-Cost Option) on Predictors for Reward (High-Cost Option Reward), Delay (High-Cost Option Delay), and their Interaction Terms.

| Parameter | Estimate | Est. Error | 2.5% | 97.5% |
|---|---|---|---|---|
| **(Intercept)** | 25.867 | 4.242 | 17.793 | 34.506 |
| **Reward** | 72.914 | 11.447 | 50.953 | 96.234 |
| **Delay** | -3.793 | 1.036 | -5.985 | -1.853 |
| **Reward x Delay** | -6.699 | 2.927 | -12.868 | -1.206 |

**Supplementary Table 6.** Bayesian Linear Mixed Models of the Effort Discounting Task, Regressing Decision Times on Predictors for Drug, Reward (Difference between High-Cost vs. Low-Cost Reward Level), Effort (Difference between High-Cost vs. Low-Cost Effort Level), and their Interaction Terms.

| Parameter | Estimate | Est. Error | 2.5% | 97.5% |
|---|---|---|---|---|
| **(Intercept)** | 7.047 | 0.038 | 6.973 | 7.124 |
| **Biperiden** | -0.009 | 0.027 | -0.062 | 0.045 |
| **Haloperidol** | 0.016 | 0.032 | -0.047 | 0.078 |
| **Reward** | -0.183 | 0.015 | -0.213 | -0.154 |
| **Effort** | 0.084 | 0.011 | 0.063 | 0.105 |
| **Biperiden x Reward** | 0.005 | 0.015 | -0.024 | 0.035 |
| **Haloperidol x Reward** | 0.036 | 0.014 | 0.009 | 0.063 |
| **Biperiden x Effort** | 0.002 | 0.012 | -0.022 | 0.025 |
| **Haloperidol x Effort** | -0.021 | 0.013 | -0.047 | 0.003 |
| **Reward x Effort** | 0.083 | 0.016 | 0.051 | 0.116 |
| **Biperiden x Reward x Effort** | -0.020 | 0.023 | -0.067 | 0.026 |
| **Haloperidol x Reward x Effort** | 0.000 | 0.023 | -0.045 | 0.047 |

**Supplementary Table 7.** Bayesian Linear Mixed Models of the Delay Discounting Task, Regressing Decision Times on Predictors for Drug, Reward (High-Cost Option Reward), Delay (High-Cost Option Delay), and their Interaction Terms.

| Parameter | Estimate | Est. Error | 2.5% | 97.5% |
|---|---|---|---|---|
| **(Intercept)** | 6.974 | 0.028 | 6.922 | 7.032 |
| **Biperiden** | 0.019 | 0.023 | -0.027 | 0.065 |
| **Haloperidol** | -0.077 | 0.023 | -0.123 | -0.031 |
| **Reward** | -0.168 | 0.011 | -0.190 | -0.145 |
| **Delay** | 0.036 | 0.008 | 0.020 | 0.053 |
| **Biperiden x Reward** | 0.002 | 0.010 | -0.017 | 0.022 |
| **Haloperidol x Reward** | 0.029 | 0.009 | 0.011 | 0.046 |
| **Biperiden x Delay** | -0.003 | 0.008 | -0.020 | 0.013 |
| **Haloperidol x Delay** | -0.020 | 0.008 | -0.036 | -0.003 |
| **Reward x Delay** | 0.005 | 0.011 | -0.017 | 0.027 |
| **Biperiden x Reward x Delay** | 0.013 | 0.016 | -0.019 | 0.044 |
| **Haloperidol x Reward x Delay** | -0.002 | 0.016 | -0.033 | 0.030 |

**Supplementary Table 8.** Fixed effects from robust linear regression model with $\kappa$ as dependent variable and questionnaire total scores, sex, and age as independent variable for the effort discounting task.

| Variables | Parameter Estimates | Standard Error | *z* | *p* |
|---|---|---|---|---|
| **(Intercept)** | -0.059 | 0.051 | -1.157 | 0.252 |
| **Sex** | 0.010 | 0.016 | 0.661 | 0.511 |
| **Age** | 0.007 | 0.002 | 2.770 | **0.008** |
| **BIS-15** | -0.006 | 0.008 | -0.828 | 0.411 |
| **AES** | 0.006 | 0.009 | 0.709 | 0.481 |
| **BDI** | -0.006 | 0.011 | -0.584 | 0.562 |

**Supplementary Table 9.** Fixed effects from robust linear regression model with $\kappa$ as dependent variable and questionnaire total scores, sex, and age as independent variable for the delay discounting task.

| Variables | Parameter Estimates | Standard Error | *z* | *p* |
|---|---|---|---|---|
| **(Intercept)** | -4.621 | 1.645 | -2.808 | **0.007** |
| **Sex** | -0.510 | 0.531 | -0.962 | 0.340 |
| **Age** | 0.011 | 0.076 | 0.146 | 0.885 |
| **BIS-15** | -0.197 | 0.282 | -0.700 | 0.487 |
| **AES** | 0.458 | 0.312 | 1.471 | 0.147 |
| **BDI** | -0.023 | 0.217 | -0.108 | 0.915 |

**Supplementary Table 10.** Fixed effects from robust linear regression model with $\kappa$ as dependent variable and questionnaire subscales as independent variable for the effort discounting task.

| **Variables** | **Parameter Estimates** | **Standard Error** | ***z*** | ***p*** |
|---|---|---|---|---|
| **(Intercept)** | 0.098 | 0.007 | 13.088 | **< 0.001** |
| **BIS-15 attentional** | 0.005 | 0.011 | 0.497 | 0.6212 |
| **BIS-15 motor** | -0.025 | 0.013 | -1.863 | 0.0678 |
| **BIS-15 non-planning** | 0.010 | 0.007 | 1.475 | 0.1459 |
| **AES - Apathy** | 0.016 | 0.009 | 1.781 | 0.0804 |
| **AES - Disinterest** | -0.008 | 0.007 | -1.083 | 0.2835 |
| **AES – Social Withdrawal** | 0.007 | 0.011 | 0.579 | 0.5650 |

**Supplementary Table 11.** Fixed effects from robust linear regression model with $\kappa$ as dependent variable and questionnaire subscales as independent variable for the delay discounting task.

| Variables | Parameter Estimates | Standard Error | *z* | *p* |
|---|---|---|---|---|
| **(Intercept)** | -4.728 | 0.228 | -20.717 | **< 0.001** |
| **BIS-15 attentional** | -0.605 | 0.336 | -1.801 | 0.0771 |
| **BIS-15 motor** | 0.039 | 0.345 | 0.114 | 0.9100 |
| **BIS-15 non-planning** | 0.224 | 0.286 | 0.783 | 0.4372 |
| **AES - Apathy** | 0.311 | 0.294 | 1.056 | 0.2957 |
| **AES - Disinterest** | 0.432 | 0.262 | 1.647 | 0.1054 |
| **AES – Social Withdrawal** | -0.252 | 0.232 | -1.086 | 0.2824 |

**Supplementary Table 12.** Bayesian Generalized Linear Mixed Models of the Effort Discounting Task – Fatigue Effects; Regressing Choices (High-Cost vs. Low-Cost Option) on Predictors for Drug, Reward (Difference between High-Cost vs. Low-Cost Reward Level), Effort (Difference between High-Cost vs. Low-Cost Effort Level), and their Interaction Terms, as well as Trial Number and two-way Trial number x Drug interactions.

| Parameter | Estimate | Est. Error | 2.5% | 97.5% |
|---|---|---|---|---|
| **(Intercept)** | 2.602 | 0.224 | 2.169 | 3.056 |
| **Biperiden** | 0.606 | 0.210 | 0.206 | 1.030 |
| **Haloperidol** | -0.475 | 0.216 | -0.903 | -0.045 |
| **Reward** | 3.516 | 0.256 | 3.037 | 4.049 |
| **Delay** | -1.682 | 0.122 | -1.919 | -1.438 |
| **Trial Number** | -0.605 | 0.073 | -0.747 | -0.462 |
| **Biperiden x Reward** | 0.792 | 0.299 | 0.218 | 1.398 |
| **Haloperidol x Reward** | -0.196 | 0.272 | -0.718 | 0.348 |
| **Biperiden x Delay** | 0.012 | 0.157 | -0.294 | 0.328 |
| **Haloperidol x Delay** | 0.024 | 0.124 | -0.221 | 0.274 |
| **Biperiden x Trial Number** | 0.142 | 0.107 | -0.068 | 0.350 |
| **Haloperidol x Trial Number** | -0.431 | 0.101 | -0.632 | -0.232 |
| **Reward x Delay** | 0.132 | 0.186 | -0.241 | 0.490 |
| **Biperiden x Reward x Delay** | 0.117 | 0.298 | -0.480 | 0.676 |
| **Haloperidol x Reward x Delay** | -0.262 | 0.248 | -0.740 | 0.238 |

**Supplementary Table 13.** Bayesian Generalized Linear Mixed Models of the Delay Discounting Task – Fatigue Effects; Regressing Choices (High-Cost vs. Low-Cost Option) on Predictors for Drug, Reward (High-Cost Option Reward), Delay (High-Cost Option Delay), and their Interaction Terms, as well as Trial Number and two-way Trial number x Drug interactions.

| Parameter | Estimate | Est. Error | 2.5% | 97.5% |
|---|---|---|---|---|
| **(Intercept)** | 20.230 | 2.637 | 15.137 | 25.461 |
| **Biperiden** | -1.318 | 1.235 | -4.043 | 0.695 |
| **Haloperidol** | -0.019 | 1.080 | -2.477 | 1.921 |
| **Reward** | 57.451 | 7.123 | 43.829 | 71.553 |
| **Delay** | -2.295 | 0.537 | -3.341 | -1.240 |
| **Trial Number** | -0.204 | 0.064 | -0.326 | -0.079 |
| **Biperiden x Reward** | -3.878 | 3.407 | -11.446 | 1.564 |
| **Haloperidol x Reward** | -1.219 | 2.927 | -7.948 | 4.017 |
| **Biperiden x Delay** | 0.792 | 0.490 | -0.106 | 1.843 |
| **Haloperidol x Delay** | 1.338 | 0.557 | 0.243 | 2.481 |
| **Biperiden x Trial Number** | 0.146 | 0.088 | -0.028 | 0.320 |
| **Haloperidol x Trial Number** | -0.003 | 0.087 | -0.173 | 0.163 |
| **Reward x Delay** | -2.808 | 1.409 | -5.618 | -0.059 |
| **Biperiden x Reward x Delay** | 1.086 | 1.363 | -1.365 | 4.061 |
| **Haloperidol x Reward x Delay** | 2.369 | 1.554 | -0.687 | 5.574 |

**Supplementary Table 14.** Bayesian Generalized Linear Mixed Models of the Effort Discounting Task – Session Effects; Regressing Choices (High-Cost vs. Low-Cost Option) on Predictors for Drug, Reward (Difference between High-Cost vs. Low-Cost Reward Level), Effort (Difference between High-Cost vs. Low-Cost Effort Level), and their Interaction Terms, as well as Session and two-way Session x Drug interactions.

| Parameter | Estimate | Est. Error | 2.5% | 97.5% |
|---|---|---|---|---|
| **(Intercept)** | 2.542 | 0.213 | 2.133 | 2.966 |
| **Biperiden** | 0.594 | 0.199 | 0.217 | 1.007 |
| **Haloperidol** | -0.542 | 0.193 | -0.924 | -0.176 |
| **Reward** | 3.411 | 0.247 | 2.944 | 3.916 |
| **Delay** | -1.648 | 0.120 | -1.883 | -1.409 |
| **Session** | 0.488 | 0.183 | 0.125 | 0.852 |
| **Biperiden x Reward** | 0.783 | 0.288 | 0.253 | 1.373 |
| **Haloperidol x Reward** | -0.284 | 0.250 | -0.776 | 0.207 |
| **Biperiden x Delay** | -0.018 | 0.149 | -0.308 | 0.279 |
| **Haloperidol x Delay** | 0.092 | 0.121 | -0.142 | 0.326 |
| **Biperiden x Session** | -0.158 | 0.305 | -0.758 | 0.445 |
| **Haloperidol x Session** | -0.426 | 0.293 | -1.018 | 0.132 |
| **Reward x Delay** | 0.148 | 0.182 | -0.216 | 0.499 |
| **Biperiden x Reward x Delay** | 0.090 | 0.290 | -0.486 | 0.642 |
| **Haloperidol x Reward x Delay** | -0.222 | 0.237 | -0.685 | 0.241 |

**Supplementary Table 15.** Bayesian Generalized Linear Mixed Models of the Delay Discounting Task – Session Effects; Regressing Choices (High-Cost vs. Low-Cost Option) on Predictors for Drug, Reward (High-Cost Option Reward), Delay (High-Cost Option Delay), and their Interaction Terms, as well as Session and two-way Session x Drug interactions.

| Parameter | Estimate | Est. Error | 2.5% | 97.5% |
|---|---|---|---|---|
| **(Intercept)** | 20.491 | 2.751 | 15.007 | 25.963 |
| **Biperiden** | -1.355 | 1.260 | -4.100 | 0.616 |
| **Haloperidol** | 0.024 | 1.121 | -2.543 | 2.007 |
| **Reward** | 58.132 | 7.390 | 43.331 | 72.868 |
| **Delay** | -2.286 | 0.539 | -3.353 | -1.230 |
| **Session** | 0.400 | 0.242 | -0.062 | 0.871 |
| **Biperiden x Reward** | -3.924 | 3.464 | -11.646 | 1.420 |
| **Haloperidol x Reward** | -1.081 | 3.032 | -8.141 | 4.302 |
| **Biperiden x Delay** | 0.787 | 0.487 | -0.087 | 1.843 |
| **Haloperidol x Delay** | 1.347 | 0.567 | 0.219 | 2.495 |
| **Biperiden x Session** | -0.064 | 0.375 | -0.807 | 0.657 |
| **Haloperidol x Session** | -0.315 | 0.370 | -1.044 | 0.392 |
| **Reward x Delay** | 1.088 | 1.349 | -1.314 | 4.003 |
| **Biperiden x Reward x Delay** | 2.430 | 1.585 | -0.660 | 5.601 |
| **Haloperidol x Reward x Delay** | -2.809 | 1.429 | -5.678 | 0.008 |
